# A highly effective and time-efficient method for extracting RNA from Cyanobacteria

**DOI:** 10.1101/2025.03.16.643587

**Authors:** Bharat Kumar Majhi

**Author notes:** Corresponding author. E-mail address (BK Majhi).

## Abstract

Cyanobacteria (blue-green algae) have been widely used as model organisms in photobiochemical research, and they have recently been exploited as hosts in numerous pilot studies to produce valuable biochemicals via genetic and metabolic modifications. The optimal production of these chemicals within the cell is dependent on several factors, including the expression of desired foreign genes. The successful expression of foreign genes within the cell translates into mRNA and corresponding proteins, which ultimately produce the desired end products. Several techniques, including RNA-sequencing (RNA-Seq), quantitative reverse transcription polymerase chain reaction (RT-qPCR), microarrays, northern blotting, and droplet digital polymerase chain reaction (ddPCR), have recently been developed to study gene expression levels in cells using ribonucleic acid (RNA) as the raw material. The quality of RNA is crucial for obtaining accurate and error-free results. Cyanobacteria have thick cellular membranes and a large spectrum of secondary metabolites, which require additional processes to break the cells and isolate RNA from cellular extracts, making extraction of high-quality RNA difficult and expensive. Using *Synechocystis* sp. PCC 6803 as a model, this study developed a highly effective and time-efficient method for extracting total RNA from cyanobacteria without the use of hazardous chemicals such as phenol and chloroform and with a minimal investment. This protocol uses standard centrifugation techniques and laboratory chemicals such as citric acid, EDTA, SDS, NaCl, and tri-sodium citrate dihydrate to extract RNA from cyanobacterial cells. The results of the quantification, purity, and integrity checks show that the quality and concentration of extracted RNA are superior to the phenol-chloroform extraction methods. Furthermore, RT-qPCR results demonstrate that the extracted RNA is of good quality and suitable for downstream applications.

**Highlights:**

- Cost-effective and time-efficient RNA extraction method.
- RNA extraction from cyanobacteria without the involvement of hazardous chemicals.
- Extraction of high-quality RNA for downstream applications.

## Introduction

Cyanobacteria are oxygenic photosynthetic organisms that have been used in various studies to investigate photosynthesis and biochemical production within the cell (Agarwal et al., 2022; Betterle et al., 2020; Betterle & Melis, 2019; Chaves et al., 2017; Jensen & Leister, 2014; Majhi, 2024; Majhi & Melis, 2024; McCormick et al., 2013; Zhang et al., 2023). They have superior natural characteristics, including mixotrophic growth (Muñoz-Marín et al., 2024), uptake of exogenous DNA in conjunction with homologous recombination (Barten & Lill, 1995; Kufryk et al., 2002), and minimal growth requirements compared to other photosynthetic organisms (Singh et al., 2016), making them ideal candidates for biotechnological applications. *Synechocystis* sp. PCC 6803 is a unicellular freshwater cyanobacterium widely used as a model organism in photosynthesis research (Ikeuchi & Tabata, 2001). It has recently emerged as an efficient host for the production of a variety of valuable chemicals, including biofuel (Majhi, 2024), biopharmaceutical proteins (Betterle et al., 2020; Majhi & Melis, 2024), and bioplastic (Agarwal et al., 2022). The production of these chemicals within the cell requires effective genetic and metabolic changes. Modifying and introducing foreign genes into the genome to synthesize desired products necessitates their effective expression. RNA analysis is an efficient and reliable method for studying gene expression levels in the genome (Corchete et al., 2020). Several techniques have been developed to analyze RNA and evaluate gene expression, including quantitative polymerase chain reaction (qPCR), RNA-seq, RT-qPCR, complementary DNA (cDNA) libraries, and northern blotting (Singh et al., 2018; Stanton, 2001; Zhang et al., 2019). Therefore, quality and quantity of RNA are particularly critical for the aforementioned downstream applications.

Extracting high-quality RNA from cyanobacteria is always challenging. Cyanobacteria have tough cellular membranes (Mohr et al., 2011), which pose challenges to the currently available RNA extraction methods. Furthermore, cyanobacteria contains a variety of secondary metabolites, including polysaccharides and phenolic compounds, which hinder the effectiveness of RNA extraction methods in producing high-quality RNA (Pinto et al., 2009). Several methods, including chemical, physical, mechanical, and heat and cold treatment processes, have been successfully used to extract RNA from cyanobacteria; however, they produce low yields (Kim et al., 2006; Kim Tiam et al., 2019; Pinto et al., 2009; Singh et al., 2010). Extracting RNA with chemicals such as organic solvents and phenol has been shown to work (Cárdenas Espinosa et al., 2021; Pinto et al., 2009); however, these chemicals pose possible health hazards (Babich & Davis, 1981; Joshi & Adhikari, 2019; Rahman et al., 2022).

In addition, it takes a long time and has low yield. Physical and mechanical procedures, on the other hand, produce higher yields than chemical ones, but require more time and processes to break the cells (Kim Tiam et al., 2019). Further, it uses certain chemicals to extract RNA from broken cells. Moreover, using high temperature to disrupt the cells and extract RNA has detrimental effect on RNA structure (Kim Tiam et al., 2019). Commercial kits have been used to circumvent the aforementioned challenges while extracting huge amounts of high-quality RNA in a short period of time. Although it has a standard procedure for extracting high-quality RNA, commercially available kits are prohibitively expensive. Furthermore, the majority of commercially available RNA purification kits are not intended for organisms that contain polysaccharides and phenolic compounds, such as plants and cyanobacteria, because these compounds irreversibly bind to nucleic acids and co-precipitate them (Dos Reis Falcão et al., 2008; Greco et al., 2014; Jensen et al., 2023; Sim et al., 2013; Wang et al., 2008). A few commercial kits have been designed to extract RNA from plant tissue but not from cyanobacteria.

This study developed a novel method for extracting total RNA from cyanobacteria using simple laboratory chemicals without using hazardous chemicals. This new method is a combination of physical and chemical procedures. It is simple, quick, inexpensive, and offers a better yield. This study used the cyanobacterium *Synechocystis* sp. PCC 6803 as a model organism for RNA extraction. This study describes a highly effective and time-efficient method for extracting total RNA from cyanobacteria for a wide range of downstream applications.

## Materials and Methods

### Cyanobacterial culture

Cyanobacterial cells were inoculated in blue green-11 (BG-11) medium in a 150 mL Erlenmeyer flask and grown at 30°C under continuous white light (30 µmol photons m^-2^ s^-1^) (Pandey et al., 2023). Cells were grown until OD730 nm reached 0.8-1 and then harvested by centrifugation at 5,000 g for 10 min. The supernatant was discarded, and the pellet was resuspended in 5 mL of BG-11 medium.

### RNA extraction

#### Method 1

1 mL of cell suspension (10 µg mL^-1^ chlorophyll *a*) was transferred to a 2 mL microfuge tube and vortexed vigorously for 5 min at maximum speed. Then 300-400 µL of lysis buffer [(0.01 M EDTA, 0.132 M anhydrous citric acid, 0.068 M tri-sodium citrate dihydrate, and 0.069 M SDS) Solution was made in DEPC-treated Milli-Q water and pH adjusted to 5] was added to the cell suspension and gently mixed by pipetting up and down 5 times using a 1000 µL pipette. Further, 150 µL of precipitation buffer [(4 M NaCl, 0.033 M anhydrous citric acid, and 0.017 M tri-sodium citrate dihydrate) Solution was made in DEPC-treated Milli-Q water] was added to the sample and mixed by inversion (10 times) followed by incubation on ice for 5-7 min. Following incubation, the sample was centrifuged for 3 min at 15,000 g on a small benchtop centrifuge to pellet out the unbroken cells and membranes. The supernatant was transferred to a new microfuge tube without disturbing the pellet and centrifuged again for 4 min at 16,000 g to remove remnants. Following centrifugation, the supernatant was transferred to another clean microfuge tube and mixed with 100% isopropanol in a 1:1 ratio. Then sample was incubated for 7 min on ice, followed by 10 min at room temperature. RNA was pelleted by centrifugation at 15,000 g for 5 min. The supernatant was discarded, and the pellet was washed with 400 µL of cold 70% ethanol and centrifuged again for 5 minutes at 15,000 g. The ethanol was carefully taken out from the tube without disturbing the RNA pellet, and the pellet was air dried by inverting the tubes on a clean paper towel for 10-15 min with the lid open. The dried pellet was then gently resuspended in 40 µL of DEPC-treated Milli-Q water and centrifuged for 5 min at 16,000 g to remove impurities and remnants (**Fig. 1**). The RNA-containing supernatant was carefully transferred to a clean microfuge tube and stored at-80°C for future analysis.

**Fig. 1.**
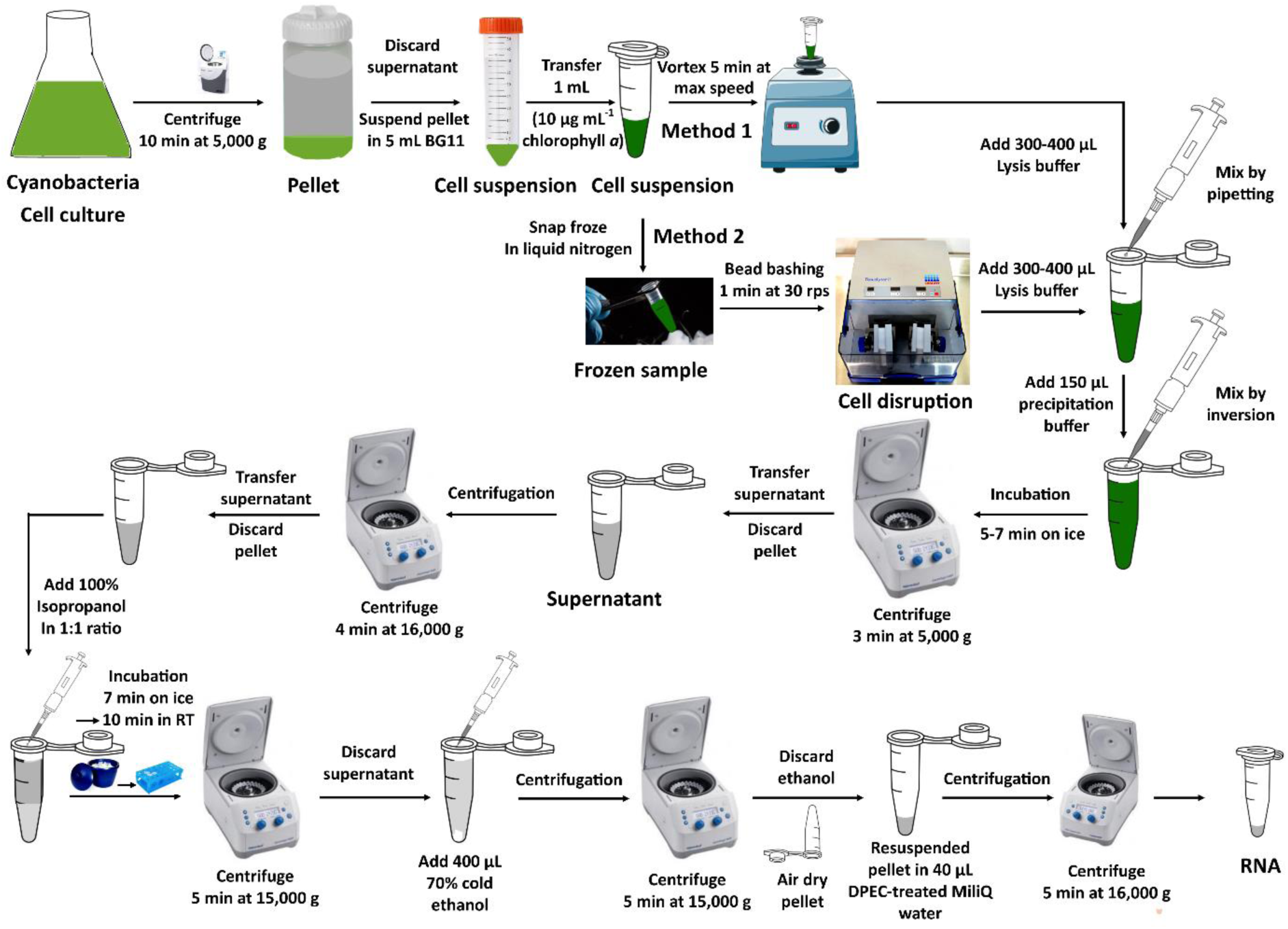
Schematic representation of cyanobacterial RNA extraction.

#### Method 2

In method 2, resuspended cells were snap frozen in liquid nitrogen and broken using stainless steel beads in a TissueLyser II (QIAGEN, catalog number: 85300) (**Fig. 1**). Frozen cyanobacterial cells were broken by bead bashing for 1 min at 30 rps. The remaining steps are the same as in method 1.

### Quantification and purity check

The quality and quantity of extracted RNA samples were assessed using nanodrop. 1 µL of the isolated RNA was used to measure the concentration and purity of RNA. The absorbance values of the isolated RNA at 260 and 280 nm (A260/280) were between 1.9 and 2.0 (**Table 1**). The concentrations of isolated RNA from methods 1 and 2 were approximately 0.125-0.150 µg/µL, and 1.2-2.0 µg/µL, respectively (**Table 1**).

**Table 1.** Concentration and absorbance values of RNA samples.

|  | **Method 1** |  | **Method 2** |  |
| --- | --- | --- | --- | --- |
| **Sample Name** | **A260/280 nm** | **Concentration**<br>**($\mu\text{g}/\mu\text{L}$)** | **A260/280 nm** | **Concentration**<br>**($\mu\text{g}/\mu\text{L}$)** |
| 1 | 2.05 | 0.125 | 1.99 | 1.53 |
| 2 | 2.0 | 0.109 | 2.04 | 1.98 |
| 3 | 1.99 | 0.150 | 2.06 | 1.89 |
| 4 | 1.95 | 0.138 | 2.03 | 1.25 |
| 5 | 1.98 | 0.132 | 1.97 | 1.77 |
| 6 | 1.99 | 0.145 | 1.95 | 1.99 |
| 7 | 1.96 | 0.149 | 1.99 | 1.36 |

### Integrity check

The quality of RNA was further evaluated on agarose gel electrophoresis. 1 µL of extracted RNA was loaded into the lanes of a 1% agarose gel and ran for 50 minutes at 150 V. The image was acquired with the BIO-RAD Gel Doc XR+ System.

### Complementary DNA (cDNA) synthesis

RNA was treated with DNase to degrade the remnant genomic DNA prior to cDNA synthesis. 1 µg of extracted RNA was incubated with DNase I (NEB #M0303S) for 30 minutes at 37°C. To verify the degradation of genomic DNA in the RNA samples, samples were run on a 1% agarose gel alongside a control. DNase-treated RNA samples were subsequently utilized to synthesize cDNA using the enzyme Superscript III reverse transcriptase (Invitrogen # 18080044) and random hexamer primers as per manufacturer’s instructions.

### Quantitative reverse transcription polymerase chain reaction (RT-qPCR)

cDNA was diluted (1:4) using DEPC-treated Milli-Q water and used to run RT-qPCR. 4 µL of diluted cDNA was mixed with 1 µL of gene-specific primers (10 µM forward and reverse primer mix) (**Table 2**) and 5 µL of 2X qPCRBIO SyGreen Blue Mix (PCRBIOSYSTEMS # PB20.17-20) to run the PCR. qPCR was run at an annealing temperature of 58°C, and the cycle number was set to 45.

**Table 2.**
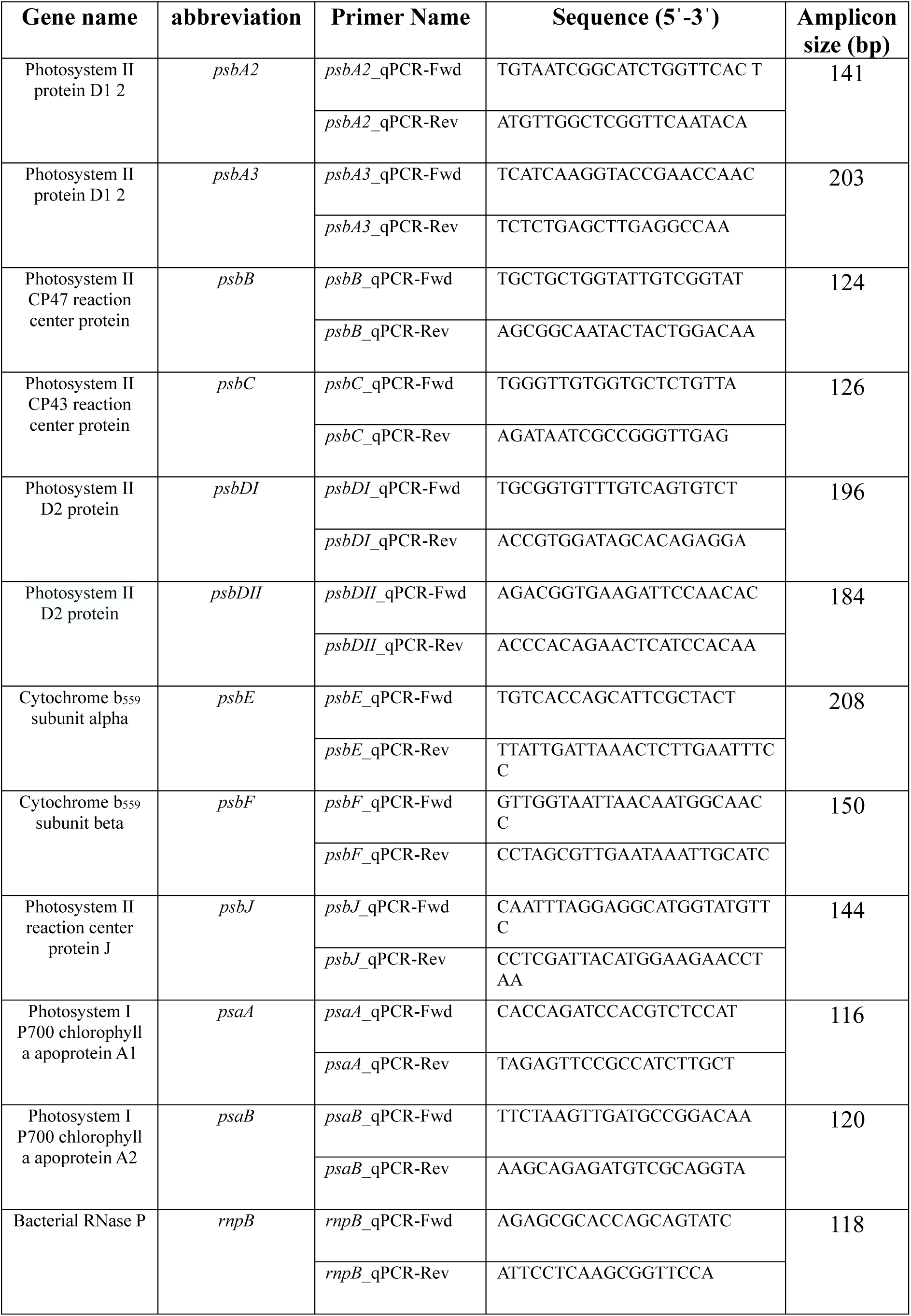
List of primers used in RT-qPCR.

## Results

### Comparative analysis between method 1 and 2

The total RNA isolated from cyanobacteria was quantified using nanodrop. The Nanodrop findings revealed substantial differences in the concentrations of RNA extracted using methods 1 and 2. Samples from method 2 yielded 1.2-2.0 µg/µL of RNA, while method 1 yielded around 0.109-0.150 µg/µL of RNA (**Fig. 2**; **Table 1**). However, the A260/280 nm values for RNA collected from both methods were similar, ranging from 1.9 to 2.0, indicating that the extracted RNA was of excellent quality.

**Fig. 2.**
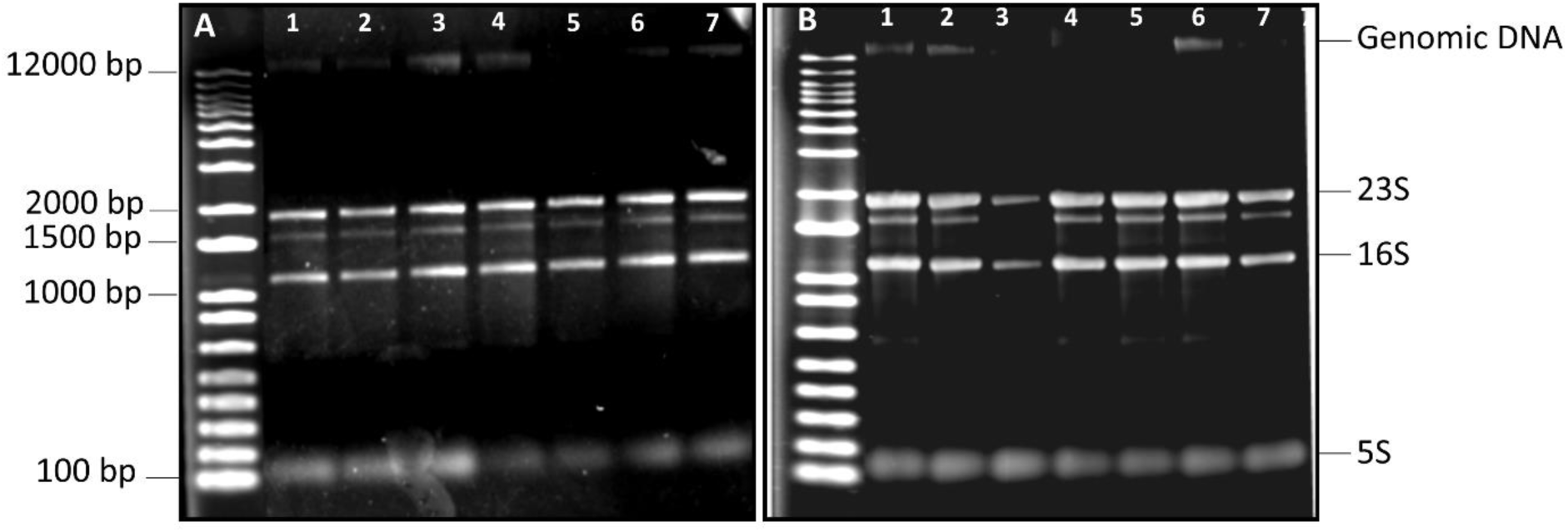
Electrophoretic mobility of extracted RNA on 1% agarose gel. (A) Total RNA extracted from samples using Method 1. (B) Total RNA extracted from samples using method 2. Samples 1-7 represent independent biological repeats.

The quality of RNA samples was further validated by performing agarose gel electrophoresis, as nanodrops have limitations in providing information about nucleic acid structural integrity. Gel electrophoresis results exhibited the presence of 23S, 16S, and 5S RNA in all of the samples (**Fig. 2**), with no degradation. The agarose gel electrophoresis results support the nanodrop result for RNA quality. In addition to RNA, all of the samples showed the presence of genomic DNA, which needs to be degraded prior to the downstream applications.

### RT-qPCR analysis

Prior to cDNA synthesis for RT-qPCR, the RNA sample was treated with DNase I to degrade the remnant genomic DNA. The DNase I treated sample was tested on a 1% agarose gel to confirm the degradation of genomic DNA (**Fig. 3A**). The gel electrophoresis result showed complete degradation of genomic DNA in the treated sample (Lane 3) compared to the untreated sample (Lane 2, lane 1 is the ladder). To further verify the complete degradation of genomic DNA in the RNA sample, PCR was performed using gene-specific primers. PCR results of the untreated sample demonstrated the presence of amplified products in lanes 2 and 4 (**Fig. 3B**); however, no PCR product was observed in the treated sample in lane 6. The absence of PCR product in the treated sample confirms the complete degradation of genomic DNA.

**Fig. 3.**
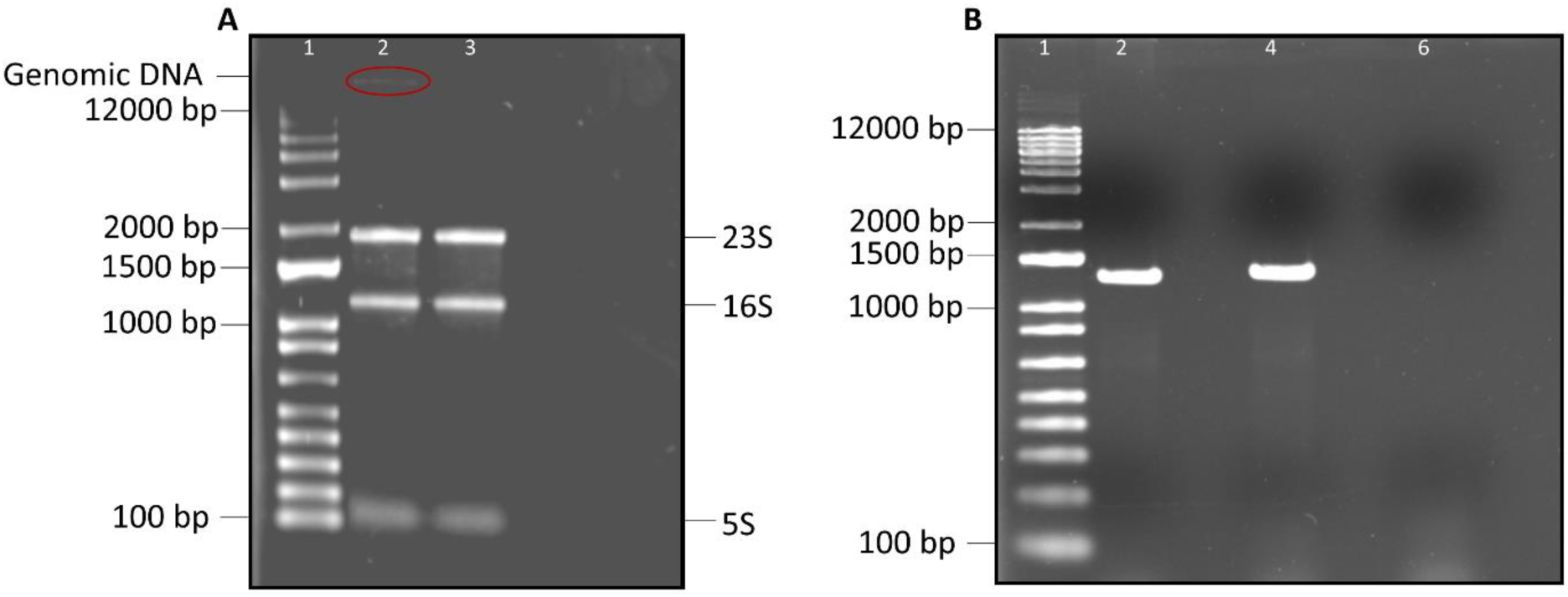
Gel electrophoresis. (A) Gel represents the electrophoretic mobility of DNase I treated and untreated RNA samples in lane 3 and 2, respectively. (B) Amplified PCR products from DNase I-untreated samples are presented in lanes 2 and 4; and treated sample in lane 6. Lane 1 represents molecular marker (1 Kb plus DNA ladder).

The DNase-treated sample was used to generate cDNA using random hexamer primers and Superscript III reverse transcriptase. To validate cDNA synthesis, samples with and without reverse transcriptase were analyzed on a 1% agarose gel. Agarose gel electrophoresis results revealed the presence of cDNA in the sample, which included the enzyme reverse transcriptase (**Fig. 4, lane 5**), indicating successful cDNA synthesis. However, no cDNA was detected in the sample, which had no reverse transcriptase (**Fig. 4, lane 3**). qPCR was performed using generated cDNA and gene-specific primers (Table 2) to investigate the gene expression levels in the genome. The qPCR findings revealed a single melting curve peak for all the genes, confirming the amplification of a single product and no primer dimer formation (**Fig. 5A**). Furthermore, the qPCR analysis exhibited steady amplification curves, confirming the quality and efficiency of the amplification (**Fig. 5B**, **Table 3**).

**Fig. 4.**
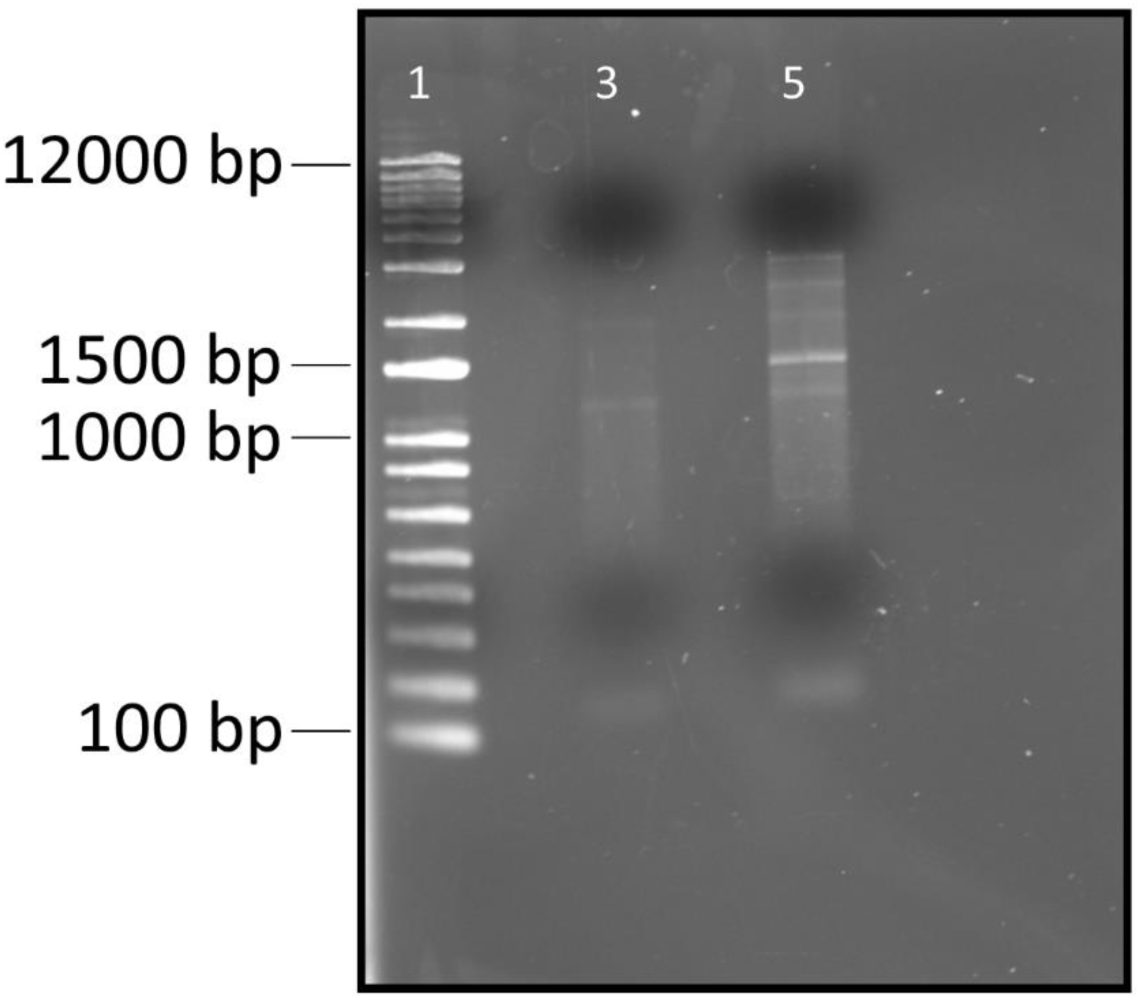
Figure represents the electrophoretic mobility of RT-PCR products (cDNA) on 1% agarose gel. Lane 1 represents 1 Kb plus DNA ladder. Lane 3 represents PCR product without Superscript III reverse transcriptase. Lane 5 represents PCR products with Superscript III reverse transcriptase.

**Fig. 5.**
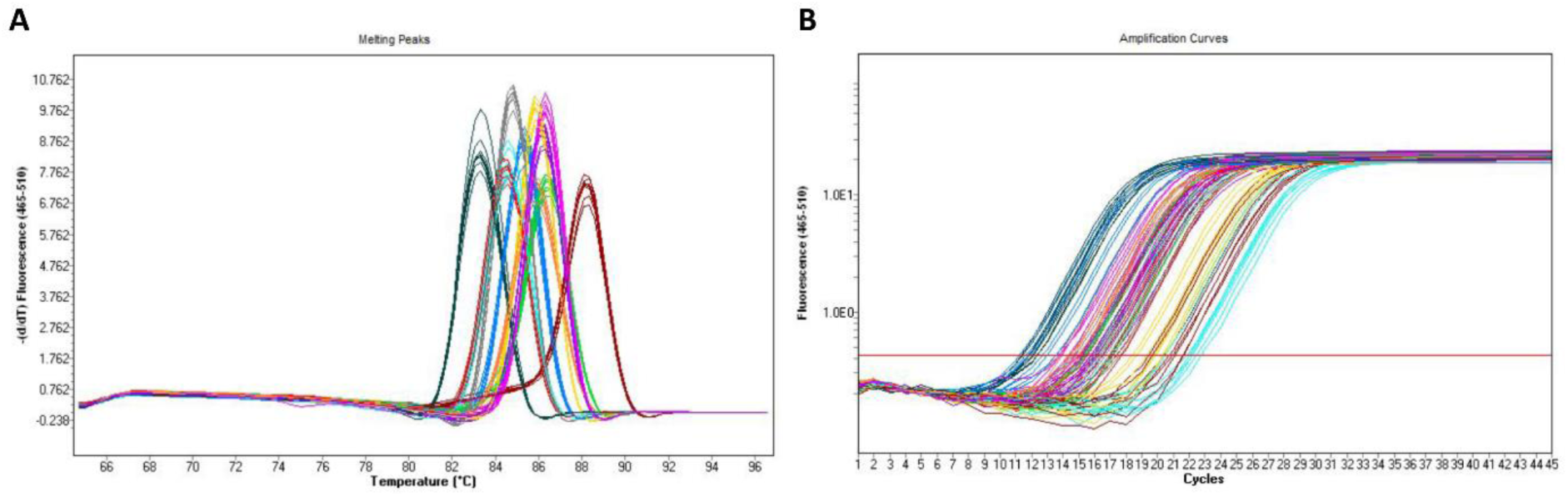
Figure represents the qPCR analysis results. Melting curves (A), and amplification curves (B).

**Table 3.** qPCR analysis results.

| Gene name | abbreviation | Amplicon T<sub>m</sub> (°C) | Ct values | Efficiency (%) |
| --- | --- | --- | --- | --- |
| Photosystem II protein D1 2 | *psbA2* | 85.31 | 15.96 | 100 |
| Photosystem II protein D1 2 | *psbA3* | 83.68 | 18.46 | 96 |
| Photosystem II CP47 reaction center protein | *psbB* | 84.48 | 17.51 | 90 |
| Photosystem II CP43 reaction center protein | *psbC* | 86.35 | 17.05 | 99 |
| Photosystem II D2 protein | *psbDI* | 86.28 | 15.92 | 89 |
| Photosystem II D2 protein | *psbDII* | 84.83 | 18.71 | 100 |
| Cytochrome b<sub>559</sub> subunit alpha | *psbE* | 85.92 | 20.81 | 99 |
| Cytochrome b<sub>559</sub> subunit beta | *psbF* | 88.15 | 21.65 | 100 |
| Photosystem II reaction center protein J | *psbJ* | 84.63 | 22.25 | 98 |
| Photosystem I P700 chlorophyll a apoprotein A1 | *psaA* | 83.35 | 12.96 | 100 |
| Photosystem I P700 chlorophyll a apoprotein A2 | *psaB* | 85.81 | 15.09 | 100 |
| Bacterial RNase P | *rnpB* | 86.22 | 18.49 | 100 |

**Table 4.** Comparison of current methods with other phenol-chloroform-based methods.

| Name of the organism | Extraction method | Concentration ($\mu\text{g}/\mu\text{L}$) | References |
| --- | --- | --- | --- |
| *Planktothrix agardhii* | Trizol based | 0.034 | (Kim Tiam et al., 2019) |
|  | Macherey-Nagel lysis buffer | 0.037-0.120 |  |
| *Anabaena variabilis* PCC 7937 | Bead-phenol-chloroform (BPC) | 0.040 | (Singh et al., 2010) |
| *Nostoc punctiforme* ATCC 29133 | PGTX | 1.29 | (Pinto et al., 2009) |
|  | Trizol | 0.47 |  |
|  | Bead-phenol-chloroform (BPC) | 0.82 |  |
| *Merismopedia tenuissima* NIES 230<br>*Synechococcus leopoliensis* UTEX 625<br>*Arthrospira platensis* NIES 39<br>*Oscillatoria tenuis* NIES 33<br>*Phormidium* sp. KCTC AG10164<br>*Synechocystis* sp. PCC 6803<br>*Anabaena flos-aquae* UTEX 2557<br>*Nodularia spumigena* UTEX 2092<br>*Nostoc* sp. PCC 7120<br>*Microcystis aeruginosa* UTEX 2666 | Bead-phenol-chloroform (BPC) | 0.202-1.327 | (Kim et al., 2006) |
|  | Xanthogenate-SDS-phenol | 0.101-0.635 |  |
|  | Lysozyme method | 0.004-0.550 |  |
| *Synechocystis* PC 6803 (This study) | TissueLyser | 1.25-2.0 |  |
|  | Vortex | 0.125-0.150 |  |

### Cost analysis

Cyanobacteria have been used by academic and industry professionals for a variety of purposes, including industrial product generation (Agarwal et al., 2022; Assunção et al., 2022; Bakku & Rakwal, 2022; do Amaral et al., 2023; Majhi, 2024; Mohanasundaram et al., 2023; Shahid et al., 2022; Velmurugan & Incharoensakdi, 2022), biopharmaceutical proteins production (Betterle et al., 2020; Majhi & Melis, 2024), and photosynthesis related research (Luan et al., 2020; Lupacchini et al., 2021; Soo et al., 2017; Treves et al., 2022; Tüllinghoff et al., 2022). The expression of heterologous genes within cyanobacterial cells is critical for the generation of the aforementioned products. To investigate genetic changes within cells, high-quality RNA or DNA is required as raw materials. Extracting high-quality RNA from cyanobacteria is always challenging due to the presence of a tough cell wall and a variety of secondary metabolites, particularly phenolic compounds. Various methods have been used to extract RNA from cyanobacteria, ranging from traditional phenol-based methods (TRIzol™ Reagent, bead-phenol-chloroform (BPC), and PGTX) to commercially available kits (Invitrogen™ PureLink™ RNA Mini Kit, RiboPure™ RNA Purification Kit, QIAwave RNA Mini Kit, Zymo Research Quick-RNA Fungal/Bacterial Microprep Kit, Attogene Plant and Algae RNA Isolation Kit, and Norgenbiotek Total RNA Purification Kits) (Pinto et al., 2009) (https://www.thermofisher.com/us/en/home/life-science/dna-rna-purification-analysis/rna-extraction/rna-types/total-rna-extraction.html; accessed 21 January 2025). Phenol-based extraction methods are more affordable than commercially available kits. Although phenol-based methods are less expensive, they do pose some health risks due to the hazardous nature of phenol. However, all of the necessary chemicals and purification kits are extremely expensive, overshadowing the overall cost-effectiveness of the process. The estimated cost for RNA extraction per cyanobacterial sample using the aforementioned chemical reagents and kits ranges between $3 and $13. (USD) (**Fig. 6**). However, the cost per sample using this new method is less than $1 (USD), which is eight to forty times less than the other methods.

**Fig. 6.**
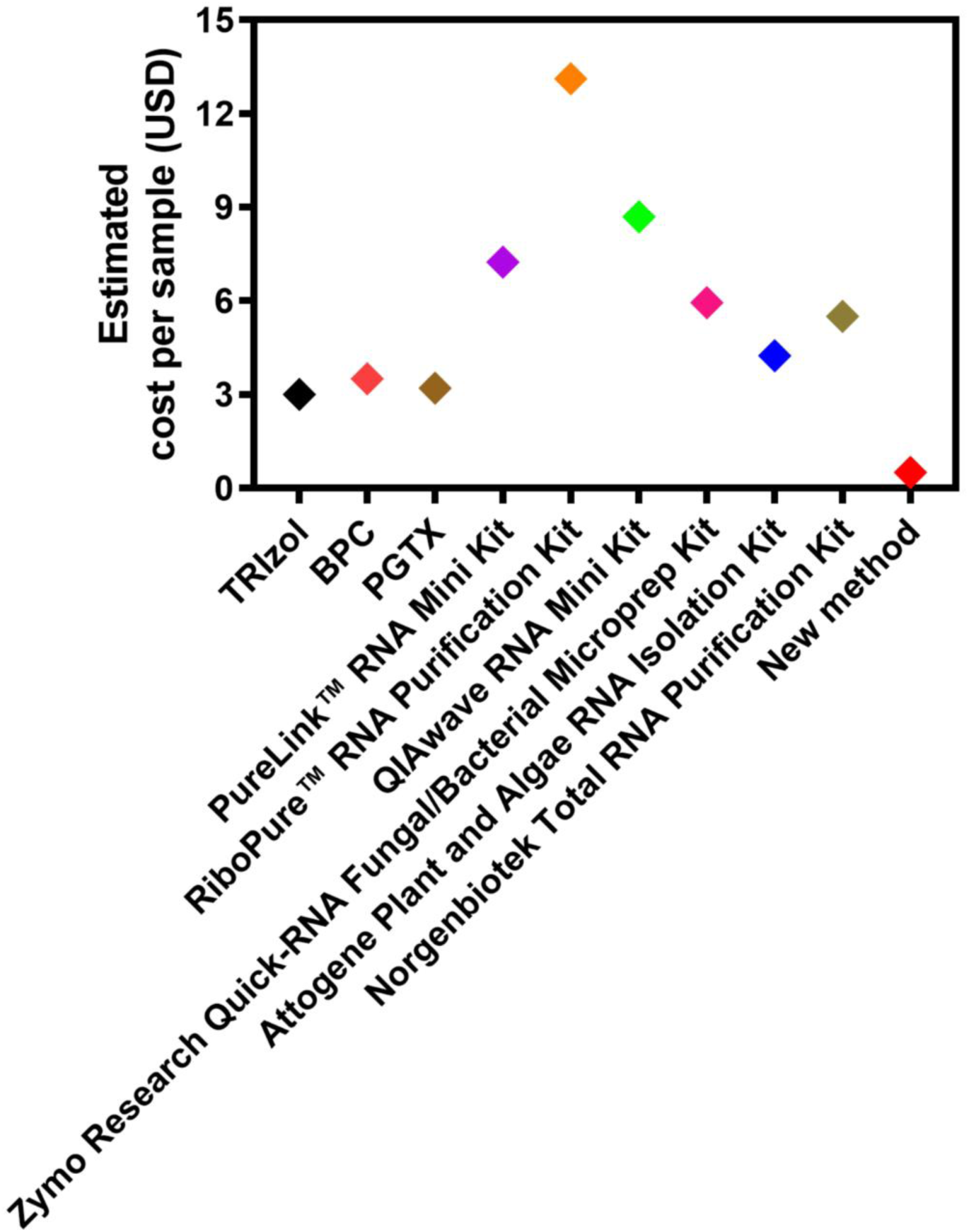
Cost comparisons of various RNA extraction methods. The cost of RNA extraction per sample is plotted along the Y-axis. Various RNA extraction methods are plotted on the X-axis. RNA extraction methods are represented in colored diamonds. Each colored diamond represent a particular RNA extraction method.

## Discussion

Cyanobacteria are the photoautotrophic prokaryotes composed of an outer cell wall and an inner cytoplasm that contains genetic material and photosynthetic apparatus (Mehdizadeh Allaf & Peerhossaini, 2022). Cyanobacterial cell wall is multilayered, consisting of a peptidoglycan layer, an outer membrane, and a surface layer (Hoiczyk & Hansel, 2000). This multilayered cell wall protects them from harsh environmental conditions. However, the thick cell wall in cyanobacteria makes RNA extraction challenging. As a result, the majority of RNA extraction methods, including phenol-based and commercial kits use a TissueLyser or vortex to homogenize the cells (Kim Tiam et al., 2019; Pathak & Lochab, 2010; Pinto et al., 2009; Yu et al., 2019). The current study also employed physical force to break the cells, using a vortex and a TissueLyser. However, the new method particularly method 2 yields more RNA than any phenol-based method (**Fig. 2B**; **Table 1** and **4**) (Kim et al., 2006; Kim Tiam et al., 2019; Pinto et al., 2009; Singh et al., 2010). Furthermore, the amount of RNA yielded by this method is comparable higher than commercial kits (**Table 5**) (Kim Tiam et al., 2019; Rump et al., 2010; Singh et al., 2010). However, the yields of RNA extracted using method 1 were comparable lower than method 2 (**Fig. 2A**; **Table 1**). It is possible that the vortexing in method 1 did not completely break down the cyanobacterial cell walls, resulting in lower yields. In addition, some phenol-based methods used heat stress rather than physical stress to disrupt cell walls and extract RNA (Kim Tiam et al., 2019; Pinto et al., 2009). Using high-temperatures (95°C-98°C) for RNA extraction may negatively impact RNA structure and quality. This novel RNA extraction method does not require any heat treatment, which can degrade RNA quality. Furthermore, unlike other phenol-based methods, this method did not use any hazardous chemicals such as phenol and chloroform, which can cause health problems (Kim Tiam et al., 2019; Pinto et al., 2009). So this method is completely safe to use.

**Table 5.** Comparison of current methods with commercially available kits.

| Name of the organism | Extraction method | Concentration ($\mu\text{g}/\mu\text{L}$) | References |
| --- | --- | --- | --- |
| *Planktothrix agardhii* | Macherey-Nagel NucleoSpin RNA, Mini kit | 0.037-0.120 | (Kim Tiam et al., 2019) |
| *Anabaena variabilis* PCC 7937 | RiboPure™-Bacteria Kit | 0.145 | (Singh et al., 2010) |
| *Salmonella enterica* | RNeasy Mini Kit | 0.78 | (Rump et al., 2010) |
|  | PureLink RNA Mini Kit | 0.97 |  |
|  | RiboPure-Bacteria Kit | 0.28 |  |
|  | UltraClean Microbial RNA Isolation Kit | 0.05 |  |
| *Synechocystis* PC 6803 (This study) | TissueLyser | 1.25-2.0 |  |
|  | Vortex | 0.125-0.150 |  |

The quality and purity of RNA are extremely important for downstream applications. A 260/280 nm ratio of 2.0 is generally considered pure for RNA (Joseph, 2016). It has been observed that RNA extracted using some of the phenol-based methods such as Trizol and BPC had a low A260/280 ratio (Pinto et al., 2009). It is possible that the RNA sample was contaminated with phenol or proteins. Compared to other methods, all the RNA extracted using the current method had a 260/280 ratio around 2.0, which confirms that all the RNA samples are free of contamination and suitable for downstream applications (**Table 1**). Although the extracted RNA had a good A260/280 ratio, all of the samples had minimal genomic DNA contamination (**Fig. 2**). The concentration of genomic DNA in RNA samples ranges from 1 to 5%. Similar results were also observed in phenol-based RNA extraction methods (Pinto et al., 2009). However, the majority of RNA extracted using kits is free of genomic DNA contamination because DNase is used to degrade genomic DNA during the extraction process. Moreover, most of the commercially available RNA extraction kits are designed for mammalian cells, plants, algae, bacteria, and yeasts. Cyanobacteria-specific commercial kits are currently unavailable.

Furthermore, the integrity of RNA is critical for subsequent applications. Integrity measurements are done using a variety of methods. The majority of studies use agarose gel electrophoresis to determine the integrity of RNA. However, the Bioanalyzer (Agilent Technologies, Inc., CA, USA) has recently been used to check the integrity of RNA (Schroeder et al., 2006). The current study used the gel electrophoresis method to ensure the integrity of extracted RNA. The agarose gel electrophoresis results exhibited clear distinct bands corresponding to 23S, 16S, and 5S RNA with no degradation, confirming the integrity of RNA (**Fig. 2**). In addition, the extracted RNA was used to perform downstream applications such as RT-qPCR to determine whether the RNA extracted by this novel method can be used to study the gene expression levels in cyanobacteria. RT-qPCR is one of the most effective techniques for studying gene expression levels in the genome.

Prior to performing RT-qPCR, the RNA sample was treated with DNase to remove genomic DNA as the sample had genomic DNA contamination (**Fig. 3**). cDNA was successfully synthesized from a DNase-treated sample using random hexamers and the enzyme Superscript III reverse transcriptase (**Fig. 4; lane 5**). The resulting cDNA was used to perform qPCR to investigate the expression levels of various genes in the *Synechocystis* sp. PCC 6803 genome. The qPCR analysis revealed a single melting curve peak for each gene (**Fig. 5A**). The absence of multiple peaks confirmed the amplification of a single product, with no non-specific amplification (Ma et al., 2021). Non-specific amplification can lead to inaccurate results. The efficiency of the qPCR assay in the range of 90% to 110% is generally considered good (Harshitha & Arunraj, 2021). The results in Table 3 show that the efficiency of the qPCR assay for all tested genes ranges between 89% and 100%, confirming the quality of the RNA and the robustness of the assay. The aforementioned qPCR analysis indicates that the RNA obtained by this method is of high quality and can be used as a raw material for downstream applications. Moreover, this current RNA extraction method is very cost-effective than other methods (**Fig. 6**).

The results presented above demonstrate the efficiency and effectiveness of this method for extracting cyanobacterial RNA. Although RNA extracted from cyanobacteria using this method contains very little genomic DNA, it can be successfully degraded by DNase treatment before being used in downstream applications. However, further optimization is required to avoid genomic DNA contamination. In addition, no hazardous chemicals are used in this method, which may cause health issues. Moreover, this method is both cost-effective and time-efficient; it can be completed in less than an hour. This method can also be used to extract RNA from other cyanobacteria, as well as plant tissues.

## Acknowledgement

I would like to thank Professor Julian Eaton-Rye at the University of Otago, New Zealand, for providing resources for this work.

## Contributions

B. K. Majhi: original concept, culture experiments, data analysis, drafting and editing manuscript.

